# A comparison of BirdNET, expert listening and acoustic indices to monitor avian diversity in a Mediterranean agricultural landscape

**DOI:** 10.64898/2026.05.20.726349

**Authors:** Irmak Akoğlu, Ergün Bacak, Sercan Bilgin, Muratcan Duran, Kerem Ali Boyla, Çağlar Akçay, Pinar Ertör-Akyazi

## Abstract

Passive acoustic monitoring poses an immense potential to assess avian diversity in many habitats, including agricultural landscapes. At the same time, automated recorders generate large datasets which present a challenge for processing and effectively assessing biodiversity. Methods such as manual listening by experts, automated detection algorithms like BirdNET and calculating acoustic indices all present different trade-offs in assessment of biodiversity through passive acoustic monitoring. In the present study we recorded soundscapes in a low-intensity agricultural landscape in western Türkiye in all four seasons. Two expert ornithologists listened to a subset of these recordings identifying bird species from the recordings. We also ran the same sample of recordings on BirdNET to compare BirdNET detections with expert detections and calculated acoustic indices for each recording. The results showed that BirdNET detected more species than experts, although some may not be reliable detections. Two acoustic indices (bioacoustic index and acoustic complexity index) were correlated positively with number of species detected by experts and one (normalized difference soundscape index) with number of species detected by BirdNET but the correlations were modest. The results show that acoustic indices may have limited value in detecting biodiversity and automated detection algorithms may do a better job, although these may need to be trained with local data to improve detection and classification.

## 1. Introduction

Intensification of agriculture is one of the most important drivers of biodiversity loss worldwide (Leroy et al. 2026; Chamberlain et al., 2000). Over the last four decades, intensive agricultural practices (e.g. monocultures, landscape homogenisation, excessive use of pesticides) have reduced farmland biodiversity, including the diversity of insects, reptiles and birds (Brown et al., 2008; Busch et al., 2020; Raven & Wagner, 2021). Farmland bird species are declining steadily in Europe; between 1990 and 2023 the farmland birds index declined more strongly (42%) than the common bird index (15%) (European Environment Agency, 2025). In this context, low-intensity farming practices have received attention for the conservation of birds and other taxa (Morelli et al., 2014; Sutcliffe et al., 2015; Tscharntke et al. 2021), as these landscapes represent semi-natural habitats offering essential resources for several farmland species, including farmland birds (Doxa et al., 2010; Krüger et al., 2023; Marcacci et al., 2020; Wolff et al., 2001).

Recent studies have suggested passive acoustic monitoring (PAM) as a quick and cost-effective tool for monitoring bird diversity in agricultural lands (Remelgado et al., 2026; Biffi et al. 2024), in contrast to traditional bird surveys that are costly and time-consuming. Passive acoustic monitoring involves use of automated recording units (ARUs) which can be programmed to record continuously or with a particular recording schedule for extended periods of time, delivering a complete recording of the soundscape. These recordings can give an indication of the diversity of species present in the landscape, provided these species emit acoustic signals, as most species of animals do.

Because PAM usually generates bioacoustic data that are too large to manually listen to by experts, there has been a significant push to generate automated techniques and algorithms to quantify the biodiversity in these large recording sets. There are two broad approaches to this problem. The first is the use of algorithms relying on machine learning for automated detection and classification of acoustic events into a set of target species (Kershenbaum et al., 2025). A popular example of this is BirdNET which is widely used to detect and classify bird vocalizations (Kahl et al., 2021).

Another approach is to use acoustic indices to quantify the pattern of acoustic variation in the recordings (Alcocer et al., 2022; Buxton et al., 2018; Sueur et al., 2008). Recent studies have used the emerging field of acoustic indices for evaluating bird diversity across different land uses, including the agricultural lands (Dixon et al., 2023; Galappaththi et al., 2024; Mammides et al., 2025; Quinn et al. 2024), and for assessing the bird diversity between certified and non-certified cocoa plantations (Dröge et al., 2024), and between organic and conventional olive groves (Myers et al. 2019), for instance. There is however a growing debate whether acoustic indices can be useful indicators of biodiversity in different biomes or in any habitats at all (Sugai et al., 2026; Alcocer et al. 2022).

Türkiye is home to significant and globally important biodiversity given its geographical location which encompasses multiple biodiversity hotspots, including the Mediterranean (Şekercioğlu et al., 2011; Myers et al., 2000). However, like in the rest of the countries in the Mediterranean basin, there are severe pressures undermining farmland biodiversity, including intensification of agriculture, climate change, and land abandonment, jeopardising high conservation value (Sokos et al., 2013). Therefore, conservation efforts in the Mediterranean target the maintenance and support of the low-intensity farmlands (Morelli et al., 2014), yet, the Mediterranean low-intensity farmlands still represent an understudied region and habitat for assessing avian diversity via passive acoustic monitoring, except for a few recent studies (Myers et al., 2019; Escudero-Fuentes et al., 2026). Moreover, to the best of our knowledge, no previous study used BirdNET and acoustic indices to quantify bird diversity in Türkiye.

In this study, we examine whether avian biodiversity can be assessed through passive acoustic monitoring (PAM) in a Mediterranean low-intensity agricultural landscape in western Türkiye. We studied the utility of BirdNET and acoustic indices for monitoring bird diversity by comparing our findings against manual listening by expert ornithologists for the same PAM dataset of sound recordings collected over four seasons in 2022-2023 in the Gödence village, located in Izmir, on the western coast of Türkiye.

## 2. Methods

### Study area

The study was carried out in the farmlands of the Gödence village (38°16’11.8”N, 26°55’06.1”E), located in the Seferihisar district of the Izmir province on the western coast of Türkiye. Gödence is a mountain village at about 440 meters above sea level, well-known for its low-intensity, agroecological production of typical Mediterranean fruits such as olives, figs, grapes, pears, mastic and almonds. Seferihisar is located in the temperate zone of the Northern Hemisphere; with average annual temperature of 17.3 degrees Celsius, and average total annual precipitation of 635.6 mm (Yildirim & Asik, 2026).

In Gödence village, agricultural lands are mainly surrounded by forest patches of *Pinus brutia* (Turkish pine) and *Quercus ithaburensis* (mount tabor oak) trees, as well as typical Eastern Mediterranean scrub species such as *Quercus coccifera* (Kermes oak), *Juniperus oxcycedrus* (prickly juniper), *Nerium oleander* (oleander), *Pistacia lentiscus* (mastic), *Arbutus unedo* (wild strawberry tree/Arbutus Berry) (Gülersoy, 2014). This vegetation often co-exists with cultivated species of olives, and other fruits grown in farmlands in Gödence, offering a rich habitat for birds and other farmland and forest species.

### Soundscape recording

We recorded the soundscape using five AudioMoth devices (Open Acoustics Devices) for the five recording points (G1-G5) shown in Figure 1, for four seasons in 2022-2023 (16-bit, 48 kHz sampling rate, medium gain, no frequency filtering) in the mixed orchards of Gödence village. The orchards were selected based on the farmers’ willingness to participate in a scientific study on farmland biodiversity. Three of the recorders were placed in a 10-hectare orchard in a relatively steep area where olives, grapes, mastic/gum, figs, almonds, pears, and walnut trees are grown, and two recorders were placed in another orchard covering 5 hectares, located at a similar altitude, where younger olive trees and almond trees are grown. Both orchards are extensively managed, without any use of synthetic inputs such as pesticides and fertilizers. Dry stone walls ensure that the soil is kept intact for agricultural production, and natural irrigation ponds are used for irrigation, important landscape elements for supporting biodiversity in the Mediterranean farmlands (Beja & Alcazar, 2003; Manenti, 2014) (See Figure S1 for examples of deployment in the field).

**Figure 1.**
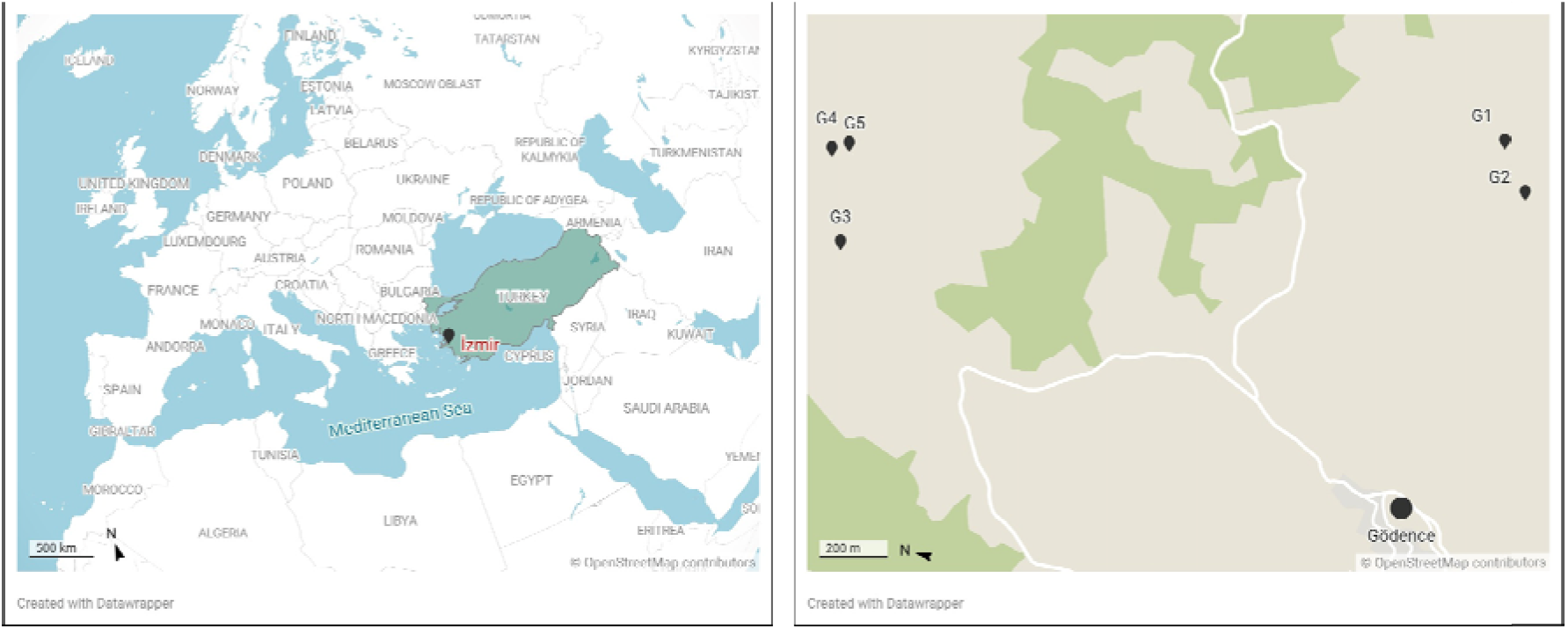
Location of Izmir, Turkey and Gödence village recording points (G1-G5) (Created with Datawrapper).

We deployed the recording devices on days without extreme weather conditions, however, as the recording devices were kept at least 5 days in the field for each season, we also obtained recordings where strong wind or heavy rain were present. We deployed the recording devices at least one meter above the ground, in trees with medium-sized trunks, in the opposite direction to the dominant wind in the region. Shaded parts of the trees were preferred to prevent overheating of the devices. Deployment to dense vegetation and running waters was avoided (Rhinehart, 2019). There were no busy roads present around the recording points. There was at least 150 meters distance between recording points in line with the general recommendation for the detection of bird vocalisations (Yip et al 2017). All recorders worked correctly as planned, however, we lost data due to memory card malfunction of one recorder (G5) in the summer season (See Figure S2 for the specific coordinates of the five recording points).

As recommended in the recent literature (Bradfer-Lawrence et al., 2019), the AudioMoth devices were programmed to record continuously for at least 120 hours per each recording point for each season (rather than subsampling on a schedule). The recordings started two hours before sunrise and continued until two hours after sunset for each day of recording. Table 1 summarizes the recording schedule and specific dates of recording employed in the study. To reduce the presence of anthropogenic sounds, we avoided the dates with labor-intensive human activity, such as harvesting and tilling periods. The recordings were made in 16-bits, 48 kHz sampling rate with medium gain setting and recorded in wave format.

**Table 1.** Recording dates and schedules for each season.

| Season | Recording Dates | Sunrise-Sunset | Total Recording Hours | Recording Hours for Each Day |
| --- | --- | --- | --- | --- |
| Fall | 8<sup>th</sup> – 13<sup>th</sup> September 2022 | 6:50 - 19:27 | 350 h | 04:30 – 21:30<br>(17 hours/day) |
| Winter | 31<sup>st</sup> January – 6<sup>th</sup> February 2023 | 8:15 - 18:36 | 467 h | 06:00 – 21:00<br>(15 hours/day) |
| Spring | 30<sup>th</sup> March – 3<sup>rd</sup> April 2023 | 6:57 - 19:34 | 308 h | 06:00 – 21:00<br>(15 hours/day) |
| Summer | 30<sup>th</sup> July – 8<sup>th</sup> August 2023 | 6:16 - 20:19 | 588 h | 05:00 – 22:00<br>(17 hours/day) |

### Manual identification of bird species richness by experts

The recording schedule described above delivered more than 1700 hours of sound data in total. The data were split into five-minute .wav files for acoustic analysis. We sampled a subset of this entire data set for further analysis by selecting the first day of each season, and the first five-minute file for each hour on that selected day. We repeated this sampling for each recording point. This process delivered 303 five-minute files in total.

Next, we manually filtered these 303 files to eliminate the silent files (files without any biophony, geophony or anthropophony), as well as the files where only non-focal sounds were present (e.g., only cicadas without any bird vocalizations) or only geophony (e.g., only heavy wind or rain without bird vocalizations). This delivered 183 five-minute files (15.25 hours in total) where bird vocalisations and some non-focal sounds of mild and moderate wind and cicadas were present. The five-minute sounds samples analysed in the present study are available upon request.

Two expert ornithologists listened manually to this subsample and identified the bird species vocalising in each five-minute file (EB and SB; each expert listened to half of the data set). The experts also noted the bird vocalisations for which they could not identify the specific species as “unidentified” bird species for each five-minute file.

### BirdNET analysis

We processed the sample (183 five-minute files) using the latest BirdNET-Analyzer v2.4, a deep learning model to identify bird sounds and species globally (Kahl et al. 2025). BirdNET analyses recordings in 3-second windows and assigns confidence scores from 0.01 to 1 for each prediction. We used the following parameters available in BirdNET: an overlap of 2 seconds (Funosas et al. 2024), detection sensitivity of 1.0 (default), and set the minimum confidence threshold at 0.1 for pre-processing the recordings. We also used BirdNET filters based on the week of the year and geographic location (Funosas et al. 2024, Perez-Granados et al. 2025)

First, the number of true positives (TP), false positives (FP), and false negatives (FN) were identified by comparing the species identified by BirdNET to the expert-identified species for each five-minute file. A detection by BirdNET was characterized as a TP if the same bird species was detected both by experts and BirdNET in the same five-minute file. It was characterized as a FP if BirdNET detected a species which was not detected by experts in the same file. A FN occurred when a species was detected in a five-minute file by the experts, but not by BirdNET. We calculated precision as an indicator for reliability of the algorithm and recall as another important indicator assessing the algorithm’s performance. Precision was calculated by dividing the TPs by the sum of TPs and FPs and recall by dividing the number of TPs by the sum of TPs and FNs. We also calculated the F1-score, which measures the predictive performance of the BirdNET algorithm by assigning equal importance to precision and recall based on the following formula:

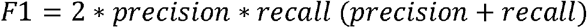

After generating predictions for each five-minute file in BirdNET with the above parameters, we identified the minimum confidence threshold that can optimize BirdNET performance for our sample based on F1-scores.

F1-score was maximized at a minimum confidence threshold of 0.27 (with an F1-score of 0.424) for our sample (See Figure S2 for the optimization of F1-score). In the rest of the analysis, we only included the species identified by BirdNET above this minimum confidence threshold of 0.27 and calculated the number of species found in BirdNET, as well as TPs, FPs, and FNs.

### Acoustic indices

We calculated five commonly used acoustic indices for our sound sample, namely, Bioacoustic Index (BIO) (Boelman et al., 2007), Normalized Difference Soundscape Index (NDSI) (Kasten et al., 2012), Acoustic Complexity Index (ACI) (Pieretti et al., 2011), Acoustic Evenness Index (AEI) and Acoustic Diversity Index (ADI) (Villanueva-Rivera et al., 2011) in Kaleidoscope Pro 5.4.8. software (Wildlife Acoustics).

We used the following parameters for the calculation of these indices: FFT Size was 512 for all indices, except for the NDSI for which 1,024 FFT Size was chosen. For the BI, 2 kHz was chosen as the minimum frequency, and 8 kHz as the maximum. For the NDSI, minimum and maximum frequencies for the anthrophony were selected as 1-2 kHz, and for biophony, as 2-8 kHz. For the ACI, the minimum frequency was taken as 0, and Nyquist frequency was chosen for the maximum frequency. The “continuous” option was selected for the number of frequency bins (denoted by J), while no dB threshold was set. Lastly, for the AEI and ADI, minimum-maximum frequencies were 0-10 kHz, and the step for frequency bands was 1 kHz. The threshold for dB was set as -50.

### Correlations between expert detections and acoustic indices

We also evaluated the performance of acoustic indices for representing bird species richness in our study area against expert detections. For this, we calculated Kendall rank correlation between the manually identified number of bird species by experts and each acoustic index for the entire dataset of the 183 five-minute sound files and corrected for multiple comparisons by taking alpha as 0.05/5= 0.01.

### Correlations between BirdNET detections and acoustic indices

Next, we assessed the performance of BirdNET against acoustic indices by calculating Kendall rank correlations between species richness detected by BirdNET and five acoustic indices for the entire dataset of the 183 five-minute sound files. Again, for each set of correlations we corrected for multiple comparisons by taking alpha as 0.05/5= 0.01.

## 3. Results

### Bird species richness via manual listening by experts

Expert listening to the 183 five-minute files delivered 36 different bird species, for a total of 784 detections in total (See Figure S3 for a summary of all the identified species by experts). On average, 4.28 ± 2.43 species were detected in each five-minute file by experts (ranging from 0 to 11). *Fringilla coelebs* (Common Chaffinch) was the most frequently detected species with presence in 123 out of the 183 files (67%), followed by other relatively abundant species of Mediterranean farmlands, including *Cyanistes caeruleus* (Eurasian Blue Tit), *Parus major* (Great Tit), *Turdus merula* (Common Blackbird), *Erithacus rubecula* (European Robin) and *Dendrocopos syriacus* (Syrian Woodpecker), present in 106, 91, 89, 61 and 40 files, respectively. There were more detections in the Spring recordings as well as at dawn and dusk (Figure S4).

### Comparing the species identified by BirdNET versus experts

For the 183 five-minute files, we compared the species identified manually by experts and by BirdNET using a minimum confidence threshold of 0.27 that optimized the F1-score. BirdNET identified 81 species in total, with a mean of 3.65 ± 2.76 species in each five-minute file (ranging from 0 to 13 for each file).

26 species were identified both by experts and BirdNET (72.2% of expert-confirmed species). 10 species were only identified by experts, but not by BirdNET: *Hirundo rustica* (Barn Swallow, detected in 12 files by experts), *Curruca cantillans* (Eastern Subalpine Warbler, detected in 5 files by experts), *Passer domesticus* (House Sparrow, detected in 4 files by experts), *Linaria cannabina* (Eurasian Linnet, detected in 2 files by experts). Six species were detected only in one file by experts, but not at all identified by BirdNET: *Luscinia megarhynchos* (Common Nightingale), *Luscinia luscinia* (Thrush Nightingale), *Dendrocopos major* (Great Spotted Woodpecker), *Galerida cristata* (Crested Lark), *Leiopicus medius* (Middle Spotted Woodpecker), and *Ficedula parva* (Red-breasted Parva).

Among the 26 species identified both by experts and BirdNET, BirdNET most frequently identified *Cyanistes caeruleus* (Eurasian Blue Tit, 987 detections), followed by *Parus major* (Great Tit, 957 detections), *Erithacus rubecula* (European Robin, 878 detections), *Turdus merula* (Eurasian Blackbird, 795 detections), and *Fringilla coelebs* (Common Chaffinch, 481 detections).

BirdNET detected 55 species that experts did not identify. Some represented plausible predictions like the *Coccothraustes coccothraustes* (Hawfinch) (45 detections) and *Muscicapa striata* (Spotted Flycatcher) (11 detections), however, there were also species that were possibly misidentified such as the *Dryobates minor* (Lesser Spotted Woodpecker),uncommon in the study area but detected 1637 times by BirdNET (more than the very common Blue Tits), which may have been the result of BirdNET confusing these calls with *Fringilla coelebs* (Common Chaffinch) calls abundant in the study area.

Table 2 provides the overall number of TPs, FPs, FNs, as well as precision and recall for our sample based on the number of total detections by BirdNET and experts in each five-minute file for a minimum confidence threshold of 0.1 versus 0.27.

**Table 2:** Assessing the performance of BirdNET via Precision, Recall and F1-score at a minimum confidence threshold of 0.10 and 0.27.

| Confidence threshold | Number of True Positives (TP) | Number of False Positives (FP) | Number of False Negatives (FN) | Precision score | Recall score | F1-score |
| --- | --- | --- | --- | --- | --- | --- |
| **0.10** | 401 | 1140 | 381 | 26.0% | 51.3% | 0.345 |
| **0.27** | 296 | 372 | 486 | 44.3% | 37.8% | 0.408 |

When we consider the subset of 23 species that were identified at least in three of the 5-minute files by experts, it is evident that increasing the minimum confidence threshold from 0.1 to 0.27 reduced the number of FPs while also slightly reducing the number of TPs, which improved the F1-score from 0.345 to 0.408 (Table 2 and Figure 2). (See Figure S6 for the same analysis of all the 36 species detected by experts).

**Figure 2.**
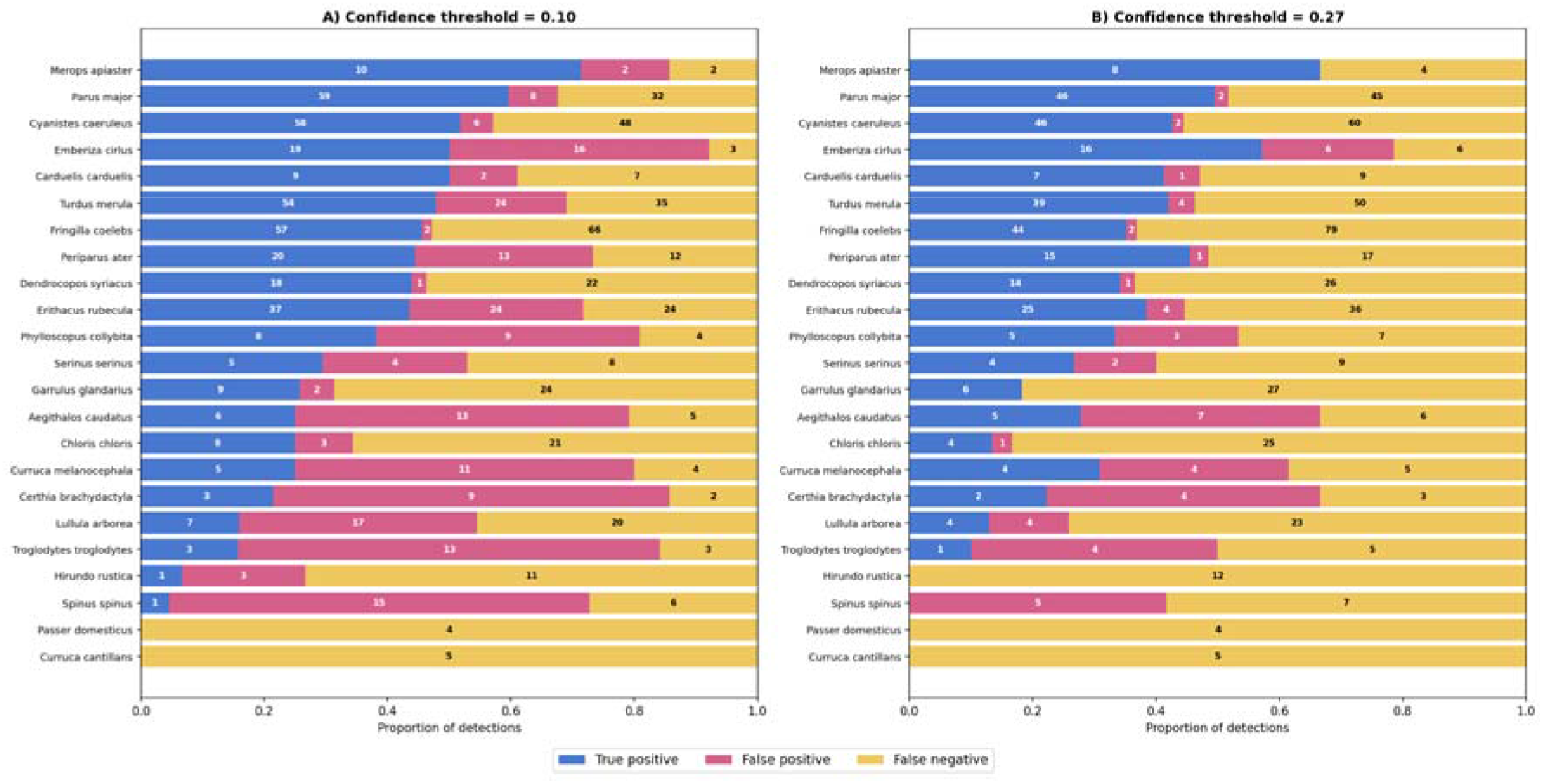
Number and proportion of TPs, FPs, and FNs for 23 bird species that were identified by experts in at least three of the five-minute files at a minimum confidence threshold of a) 0.1 and b) 0.27. The numbers within the bars represent the number of five-minute files where a TP, FP or FN occurred for each species.

### Comparing the acoustic indices with expert detections and with BirdNET detections

Next, we correlated the acoustic indices calculated from the complete sample of 183 five-minute files with the species richness detected by expert listening as well as with BirdNET detections. For the comparison between acoustic indices and expert detections, both BIO and ACI were found to be significantly positively associated with bird species richness detected by the experts after correction for multiple testing (Table 3). None of the other acoustic indices were correlated with species richness from expert listening. Looking at the correlations for the species count detected by BirdNET versus acoustic indices showed that only NDSI was significantly positively correlated with BirdNET species count after correction for multiple testing (Table 3).

**Table 3.** Kendall’s τ and associated p-values for the correlations of the five indices with species numbers detected by expert listening and BirdNET.

|  | Expert | BirdNET |
| --- | --- | --- |
| Acoustic index | Kendall’s $\tau$ (p value) | Kendall’s $\tau$ (p value) |
| BIO | **0.245 (<0.0001)** | 0.034 (0.513) |
| ACI | **0.315 (<0.0001)** | 0.115 (0.028) |
| ADI | -0.097 (0.071) | 0.034 (0.527) |
| AEI | 0.103 (0.055) | 0.033 (0.541) |
| NDSI | 0.068 (0.191) | **0.214 (<0.0001)** |

Sample size is n=183 for all cells. BIO: Bioacoustic Index, ACI: Acoustic Complexity Index, ADI: Acoustic Diversity Index, AEI: Acoustic Evenness Index, NDSI: Normalized Difference Soundscape Index. Significant correlations after correction for multiple comparisons are highlighted in bold.

## 4. Discussion

Passive acoustic monitoring is a tool with great potential for assessing avian diversity in agricultural areas. In the present study we assessed the performance of BirdNET against the expert identifications in a low-intensity Mediterranean agricultural landscape with a similar analysis adopted by Funosas et al. (2024; 2026). This was important as the growing use of BirdNET for avian monitoring needs to be evaluated in different habitats and different regions of the world. Moreover, selection of parameters in BirdNET needs to be catered to different biomes so that recall and precision can be improved. Optimising over the minimum confidence thresholds in BirdNET, we were able to increase recall and precision and improve F1-scores. Our findings showed that while there was some agreement, there were also discrepancies between the BirdNET detections and expert detections.

Next, we sought to assess the ability for five commonly used acoustic indices by correlating acoustic indices with both species detected by expert ornithologists and BirdNET. Our results showed that only BIO and ACI were positively correlated with bird species richness evaluated manually by experts. Interestingly, only NDSI was positively correlated with BirdNET species counts for the same dataset while the rest of the acoustic indices showed no significant correlation after correcting for multiple comparisons. We discuss these results in more detail below.

### Comparing expert listening and BirdNET detections

One of the main aims of our study was to evaluate BirdNET detections with expert listening. Following the approach of Funosas et al. (2024, 2026) we first optimized over the minimum confidence thresholds to maximize F1 scores which gives equal weight to both precision and recall. Interestingly, BirdNET detected many more species than the expert listeners detected. While some of these species are plausible to occur in numbers consistent with the detections, others may be influenced by False Positives (e.g. Lesser Spotted woodpecker). While expert detections are also potentially subject to error (Farmer et al. 2012), it seems likely that some, if not the majority of these detections by BirdNET are false positives.

Our findings are consistent with the recent studies that found BirdNET to be more reliable for more common species (e.g. Great Tits, Eurasian Blue Tits, Common Chaffinches). For instance, Funosas et al. (2024) found that the reliability of BirdNET detections significantly correlated with the number of Xeno-Canto recordings for that species in a European sample. We also found that increasing the minimum confidence thresholds lead to an increase in F1 scores (i.e. higher reliability), although the F1 scores remained relatively moderate.

The discrepancy between BirdNET and expert listeners is illustrated in Table 4 below with the species that are subject of special conservation measures according to the EU Directive 2009/147/EC, IUCN Red List (EU-28), and Birds@Farmland Initiative of the European Commission. The table shows that it was possible that BirdNET and experts detect the same species in different recordings, suggesting that comparison at the recording level and at the dataset level may be also useful in practice. These findings underscore the need to improve BirdNET detections in the context of conservation actions. BirdNET detections can be made more reliable with better quality recordings and potentially additional training of the model with location specific data (Funosas, 2024; 2026).

**Table 4:**
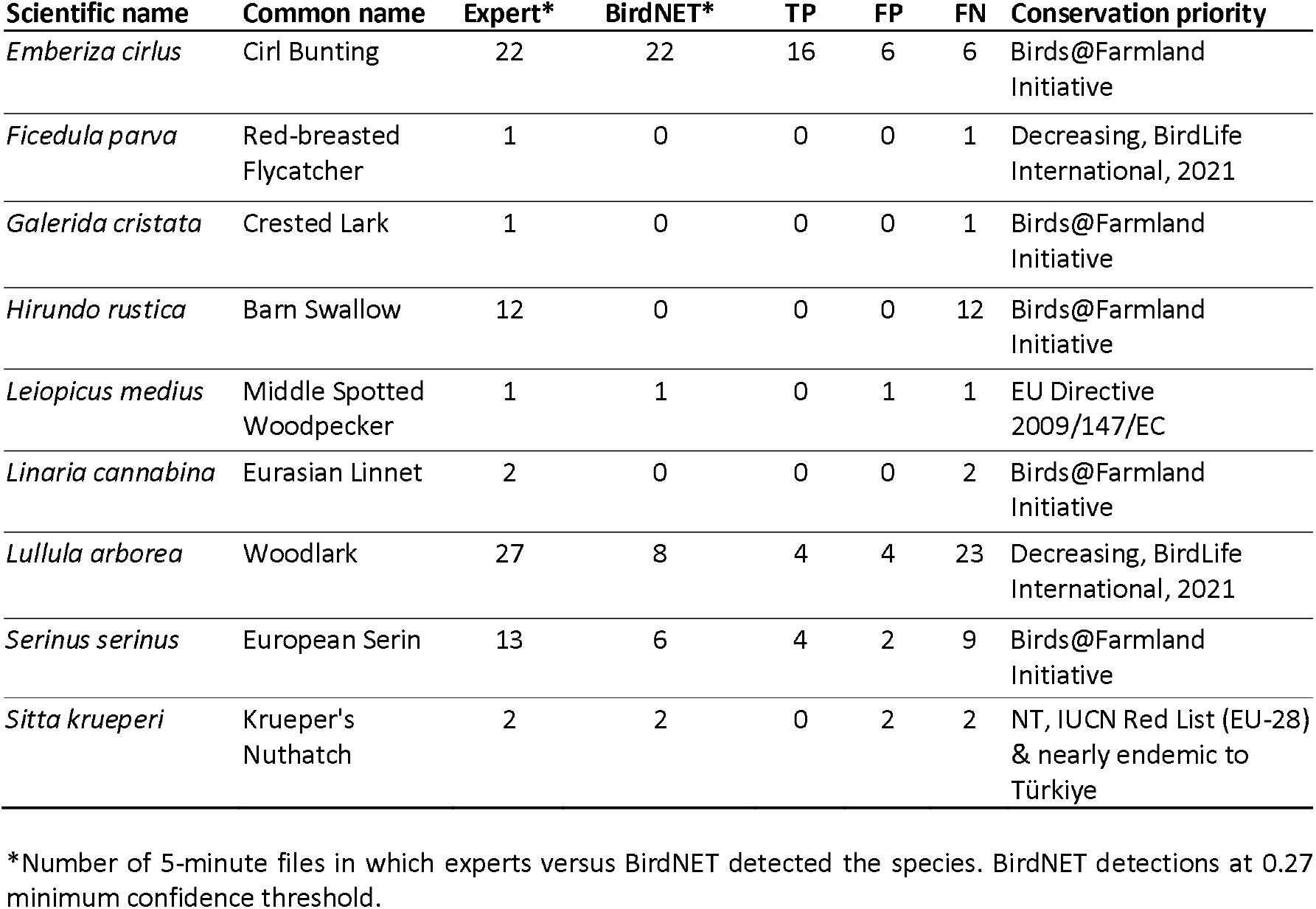
Comparison of bird species with conservation priority between experts and BirdNET identifications.

### Are acoustic indices useful in avian diversity monitoring?

Our second aim was to determine which of the acoustic indices, if any, were useful in capturing the acoustic biodiversity in our agricultural landscapes. We found that ACI and BIO were best correlated with the species richness in each recording as determined by expert listening. These results add and complement previous findings in different habitats. For instance, Budka et al. (2023) found that BIO was most strongly related to bird species richness detected by manual listening of the same sound files, whereas ACI and ADI were less strongly correlated to bird species richness. Similarly, Dröge et al. (2024) and Kotian et al. (2024) found the same relationship for BIO only, and Hilje et al. (2017) and Towsey et al (2014) for ACI only.

We also evaluated the acoustic indices with respect to another strategy of assessing biodiversity, machine learning models that detect vocalizations and classify them into species. We chose the most popular of these models, BirdNET (Kahl et al., 2021). Interestingly while BIO and ACI were the only ones correlated with species count by experts, NDSI was the only acoustic index correlated with BirdNET species detections. The superior performance of NDSI may be due to the fact that BirdNET is mostly trained with high-quality (low background noise) bird vocalisation recordings and NDSI differentiates clearly between biophony and antropophony and these will generally be well correlated (e.g. recordings with clear bird sound, i.e. biophony, will have more BirdNET detections; Funosas et al. 2024). This hypothesis is consistent with the fact that when we separated our dataset manually into categories that included clear bird calls, cicadas present, mild wind and moderate wind, NDSI performed much better in the recordings with clear bird calls (n=84 recordings) compared to the pooled sample without any sonic condition categorisation (n=183 recordings) (See Supplementary Material and Figure S5). However, further studies are to investigate this possibility in different sonic and habitat contexts.

While some acoustic indices significantly correlated with species counts by experts or BirdNET, effect sizes were rather modest. These effect sizes are consistent with the mean effect sizes in a recent meta-analysis of acoustic indices by Alcocer et al. (2022). In that meta-analysis, Alcocer and colleagues found that ACI, H (acoustic entropy index) and NDSI had the largest effect sizes in terms of correlating with biological indices of diversity. In this sense, our results are largely consistent with this overall picture (ACI being the best performing index for expert detections and NDSI the only significant index for BirdNET detections), but as they point out there are large heterogeneities among the effect sizes in the literature and indeed in our study: which index performed better depended on whether it was compared to expert listening or BirdNET detection.

Ultimately, the utility of the acoustic indices will be limited by how well they capture biological diversity and no single acoustic index may be best suited to be applied to all habitats (Galappaththi et al., 2024; Elridge et al., 2018; Mammides et al., 2025; Sethi et al. 2023). Instead, validation of the acoustic indices needs to be carried out where the acoustic indices are evaluated against biological data, e.g. bird species counts or manual listening by experts of the sound files, before widespread deployment (Alcocer et al., 2022).

Recently Sugai and colleagues (2026) argued that acoustic indices are not useful at all, given that they correlate poorly with species numbers and no explicit identification of species is possible from the acoustic indices. Instead, they argue that the current methods to automatically detect and classify sounds into species are much more effective tools.

## Conclusion

In summary, we evaluated BirdNET, expert listening and five commonly used acoustic indices in determining bird species richness in an agricultural landscape in the Eastern Mediterranean (Türkiye). Our findings suggest that while some acoustic indices correlate modestly with species numbers from expert detections, others do not and the correlations tend to have small effect sizes, potentially limiting their usefulness. BirdNET detected many more species than our experts and when they detected the same species, it was not always in the same recordings. Nevertheless, our findings suggest that passive acoustic monitoring can be a valuable tool for assessing biodiversity in the relatively understudied agricultural landscapes.

## Acknowledgements

The authors would like acknowledge Özcan Kokulu (Gödence Cooperative Head) for logistical support and arranging access to the orchards, Dr Evrim Karacetin and Dr Rasit Bilgin for their constructive criticism and discussion about the work. This research was supported by a Boğaziçi University Scientific Research Projects Start-Up Grant (No 18701) to PEA.

