## Supplementary materials for "A comparison of BirdNET, expert listening and acoustic indices to monitor avian diversity in a Mediterranean agricultural landscape"

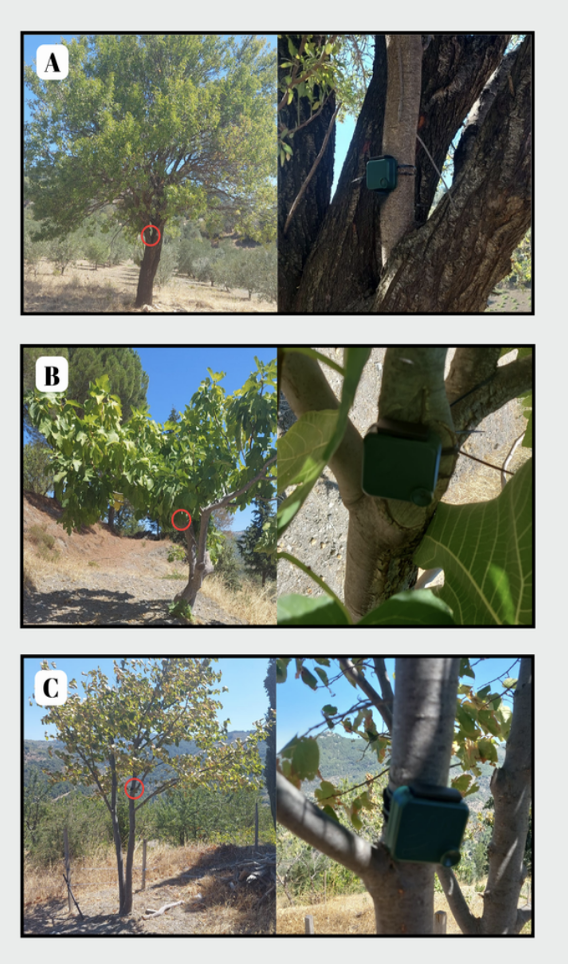

Figure S1: Deployment of AudioMoths. (A) On an almond tree at recording point G2, (B) on a fig tree at G5, (C) On a pear tree at G4

Table S1. The coordinates of the five recording points

| **Recorder ID** | **Latitude** | **Longitude** |
| --- | --- | --- |
| G1 | N 38.269822 | E 26.932366 |
| G2 | N 38.268974 | E 26.930834 |
| G3 | N 38.286129 | E 26.923265 |
| G4 | N 38.286595 | E 26.926539 |
| G5 | N 38.286995 | E 26.926262 |

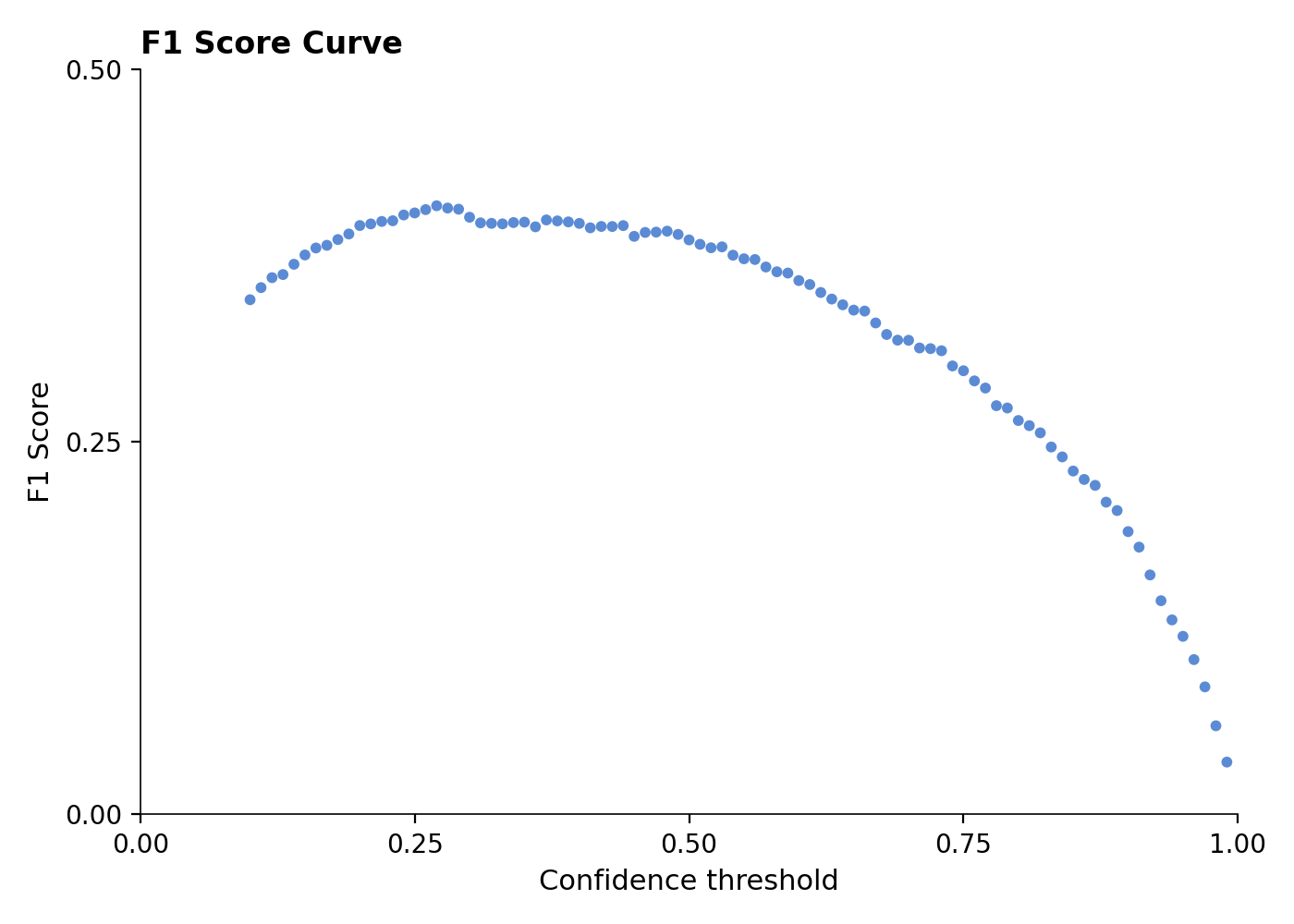

Figure S2: Optimization of the minimum confidence threshold based on the F1-score. F1-score was optimised at min. Confidence threshold of 0.27.

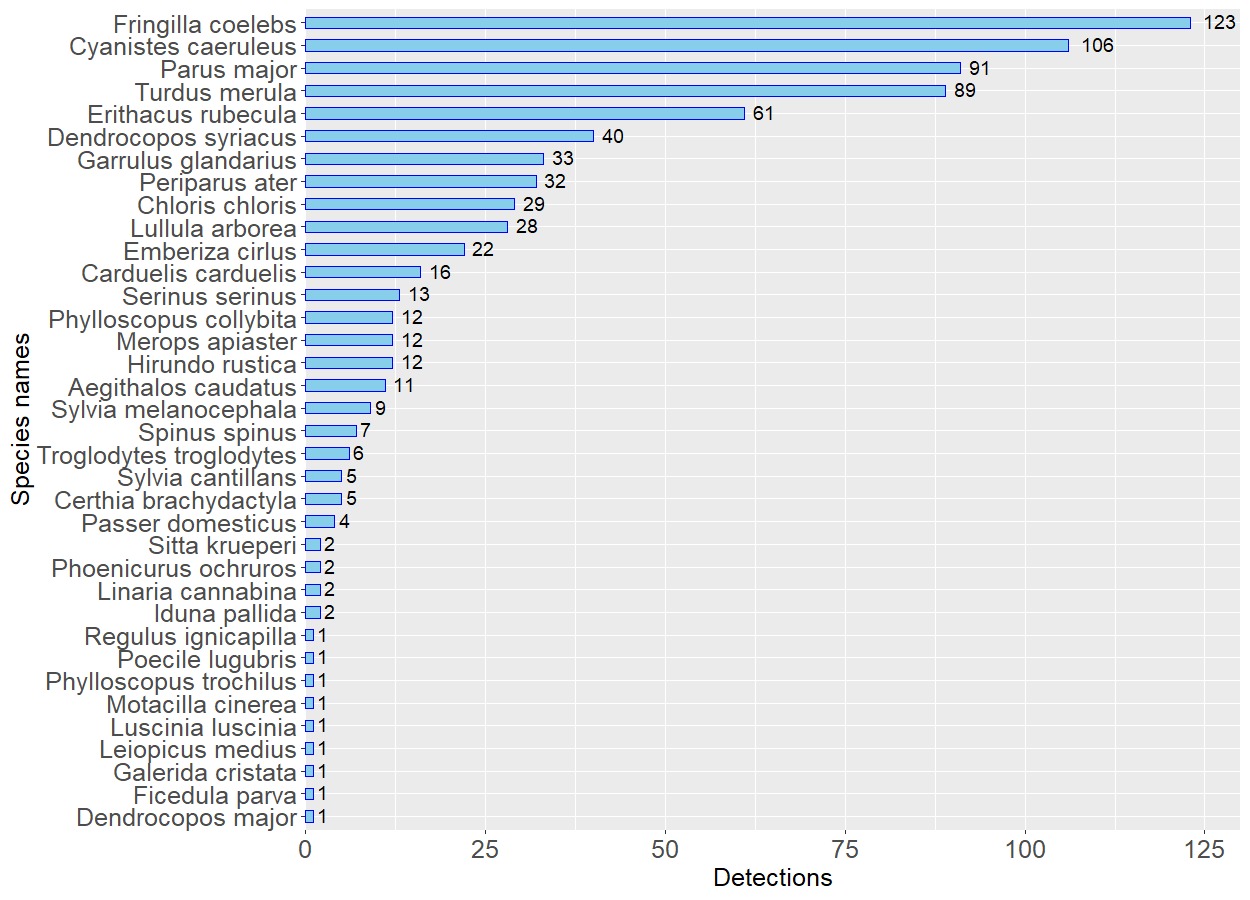

Figure S3: Bird species manually detected by experts. The numbers next to the bars indicate the total number of 5-minute files where the species has been detected by experts.

**Seasonal and daily patterns of detections by experts**

In this analysis we report the seasonal and diurnal variation in detections by experts. The number of species detected by experts for each season differed significantly, and the spring season had the highest number of species (Kruskal Wallis test with post-hoc Dunn's test using a Bonferroni correction, χ^2^(3) = 34.41, *p* < .001). Number of identified bird species changed over the day and peaked for the dawn chorus in comparison to the rest of the day (Kruskal Wallis test with post-hoc Dunn's test using a Bonferroni correction χ^2^(2) = 12.58, *p* = .002, Figure S4).

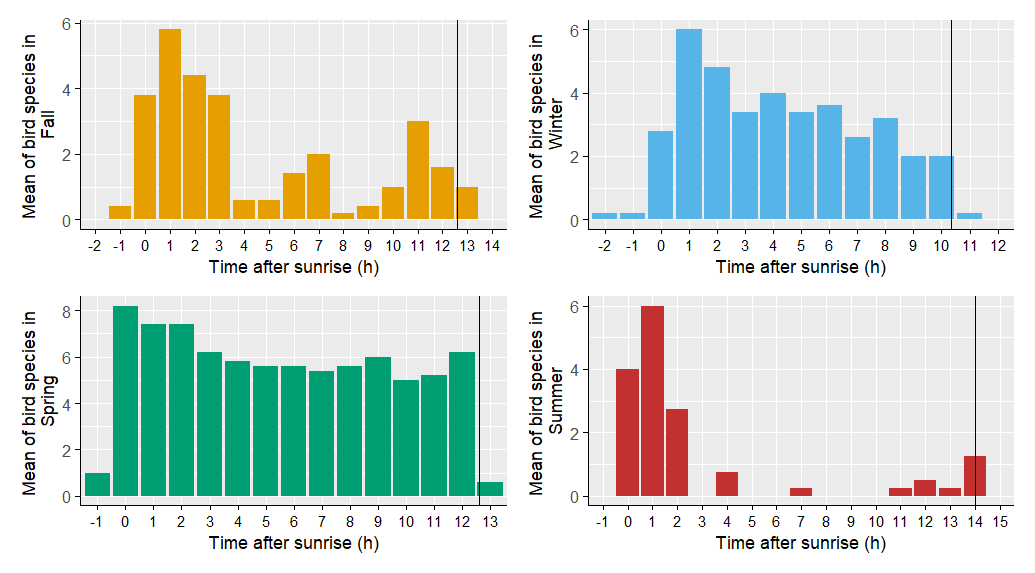

Figure S4: Hourly distribution of mean number of bird species for each season detected by experts (black vertical line representing the sunset for each season).

**Comparison of acoustic indices across seasons and times of the day**

Taking seasonality into consideration by studying the subsamples separately for each season, i.e. fall (n=46), winter (n=49), spring (n=70) and summer (n=18), we found that BIO was significantly and positively associated with bird species richness (Kendall’s *tau*=0.23, p<0.05) from expert listening in the spring season. ACI was significantly and positively associated with bird species richness (Kendall’s *tau*=0.53, p<0.05) in the summer season.

With respect to daily vocal activity, we tested the correlations between acoustic indices and bird species richness and found that for the dawn chorus (n=32), BIO (Kendall’s *tau*=0.37, p<0.001), NDSI (Kendall’s *tau*=0.35, p<0.001), and ACI (Kendall’s *tau*=0.49, p<0.001) were significantly and positively associated with bird species richness identified by experts. For the dusk chorus (n=13), BIO (Kendall’s *tau*=0.54, p<0.05), and ACI (Kendall’s *tau*=0.33, p<0.05), were significantly and positively associated with bird species richness identified by experts.

**Performance of acoustic indices in different sonic contexts**

One aim of the study was to compare the performance of the acoustic indices in different sonic contexts (Ross et al. 2021). We categorized the 183 five-minute sound files manually into four groups: (1) clear bird vocalizations without any interference from other non-focal sound sources, (2) bird vocalizations in the presence of cicadas/insects, (3) bird vocalizations in the presence of mild wind, and finally, (4) bird vocalizations under moderate wind. Seven recordings did not fall under any of these categories; therefore, they were grouped separately as “Other” (see Figure S5 for examples of spectrograms of each category). Mild wind was usually visible at below 2 kHz, thus rarely masking bird vocalizations which were typically above 2 kHz. However, moderate wind often spanned the entire spectrogram in terms of different frequencies and overlapped more strongly with bird vocalizations. When cicada vocalizations were dominant in the recordings, they were often masking the bird vocalizations, overlapping with the frequency ranges where birds vocalized.

We calculated the five acoustic indices for each sound category separately to assess the sensitivity of acoustic indices to non-focal sounds: (1) clear bird vocalizations without any interference from other sound sources (n=84), (2) bird vocalizations in the presence of cicadas/insects (n=13), (3) bird vocalizations in the presence of mild wind (n=33), and (4) bird vocalizations under moderate wind (n=46). For each of these categories, we calculated Kendall rank correlation between number of species detected by experts and acoustic indices.

For the five-minute files with clear bird vocalisations (n=84), BIO (Kendall’s *tau*=0.39, p<0.001), NDSI (Kendall’s *tau*=0.29, p<0.001), and ACI (Kendall’s *tau*=0.39, p<0.001) were significantly and positively correlated with bird species richness. In the presence of cicadas (n=13), there was a weak tendency of the ACI to be positively correlated with bird species richness (Kendall’s *tau*=0.55, p<0.05), but correlations with the rest of the acoustic indices turned out to be statistically insignificant. In the files with bird vocalisations combined with mild wind (n=33), both BIO (Kendall’s *tau*=0.47, p<0.001) and ACI (Kendall’s *tau*=0.42, p<0.001) were significantly and positively correlated with bird species richness. For the category with stronger wind (moderate wind category, n=46), none of the acoustic indices were correlated significantly with bird species richness from expert listening.

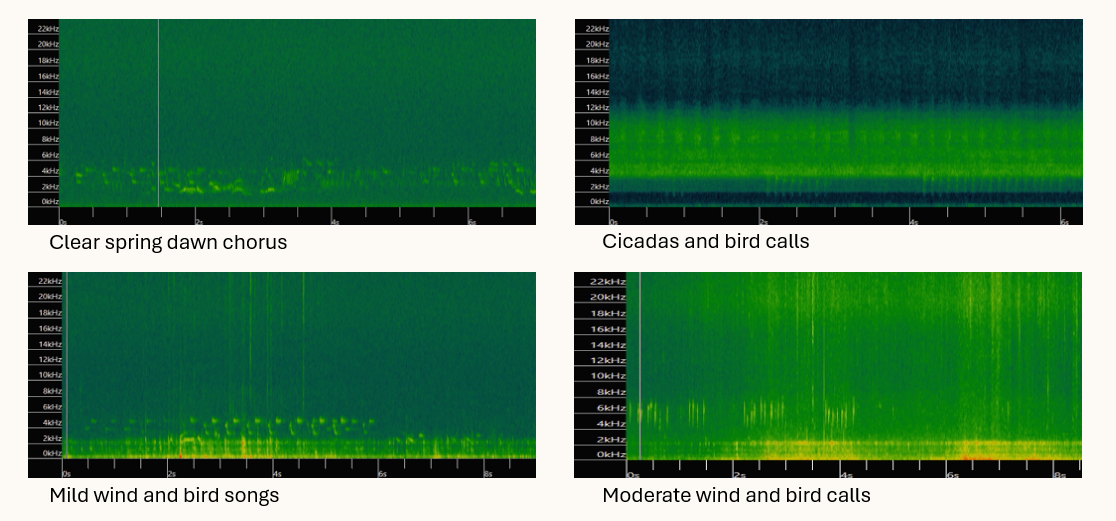

Figure S5. Examples for spectrograms of different sound categories.

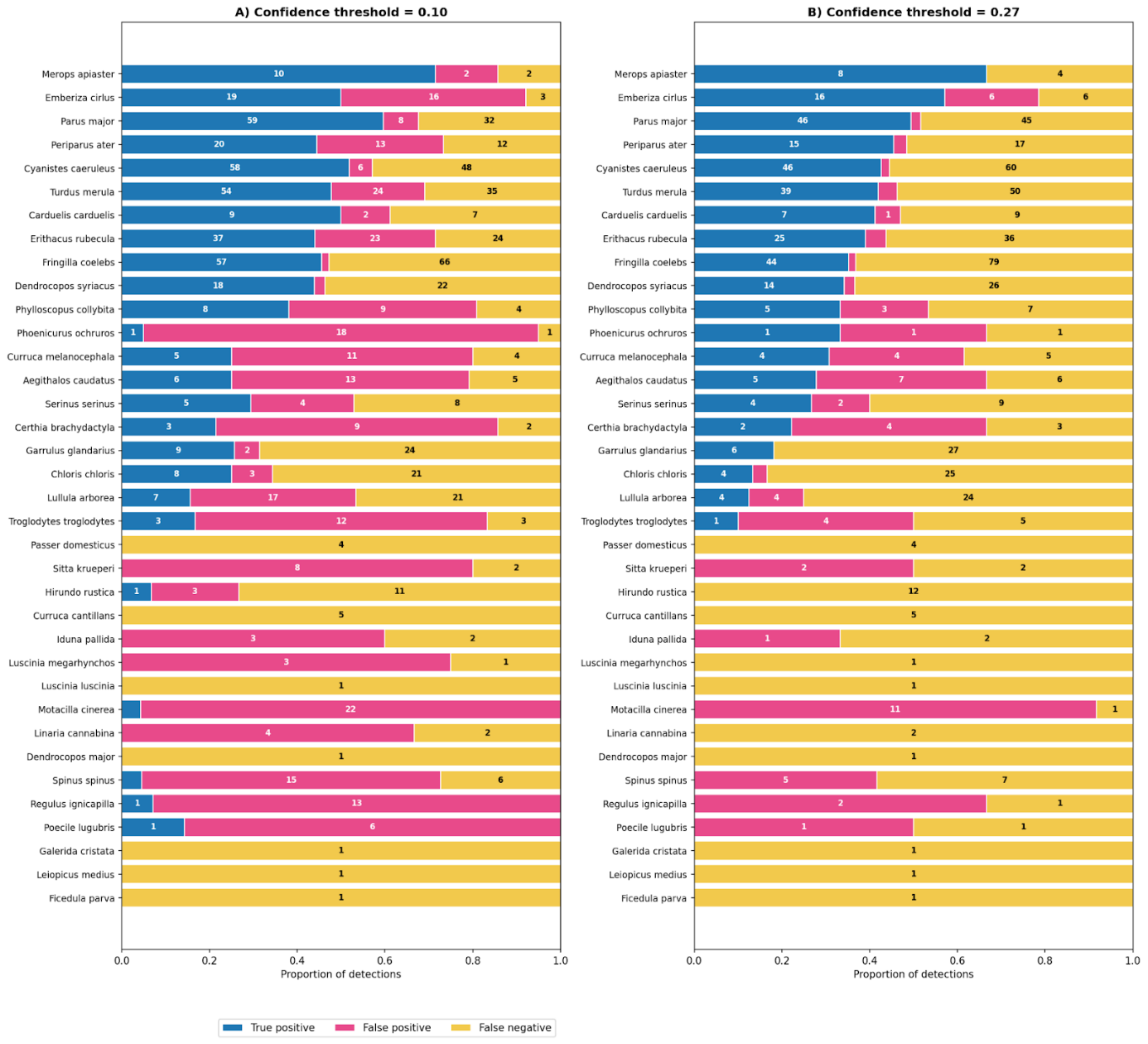

Figure S6: Number and proportion of TPs, FPs, and FNs for the total of 36 species identified by experts fat a minimum confidence threshold of a) 0.1 and b) 0.27. The numbers within the bars represent the number of five-minute files where a TP, FP or FN occurred.
